# Efficacy of lysophosphatidylcholine as direct treatment in combination with colistin against *Acinetobacter baumannii* in murine severe infections models

**DOI:** 10.1101/2020.12.02.409243

**Authors:** A Miró-Canturri, R Ayerbe-Algaba, ME Jiménez-Mejías, J Pachón, Y Smani

**Author notes:** Andrea Miró-Canturri and Rafael Ayerbe-Algaba contributed equally to this work. Author order was determined by drawing ball. **Corresponding authors**: Younes Smani, Clinic Unit of Infectious Diseases, Microbiology and Preventive Medicine, Institute of Biomedicine of Seville (IBiS), University Hospital Virgen del Rocío, Av. Manuel Siurot s/n, 41013, Seville, Spain., Manuel Enrique Jiménez-Mejías, Institute of Biomedicine of Seville (IBiS), University Hospital Virgen del Rocío, Av. Manuel Siurot s/n, 41013, Seville, Spain.

## Abstract

**Objectives:** The stimulation of the immune response to prevent the progression of the infection may be an adjuvant to antimicrobial treatment. Previously, we showed that preemptive treatment with lysophosphatidylcholine (LPC) in combination with colistin improved the therapeutic efficacy of colistin against MDR *Acinetobacter baumannii*. In this study, we aimed to evaluate the efficacy of direct treatment with LPC in combination with colistin in murine experimental models of severe infections by *A. baumannii*.

**Methods:** We used *A. baumannii* strain Ab9, which is susceptible to colistin and most of the antibiotics used in clinical settings, and *A. baumannii* strain Ab186, which is susceptible to colistin but presents a MDR pattern. The therapeutic efficacies of one and two doses of LPC (25 mg/kg/d) and colistin (20 mg/kg/8h), alone or in combination, were assessed against Ab9 and Ab186 in murine peritoneal sepsis and pneumonia models.

**Results:** One and two doses of LPC in combination with colistin and colistin monotherapy enhanced bacterial clearance of Ab9 and Ab186 from spleen, lungs and blood and reduced mortality rates compared with those of the non-treated mice group in both experimental models (*P*<0.05). Moreover, one and two doses of LPC reduced the bacterial concentration in tissues and blood in both models, and increased mice survival in peritoneal sepsis model for both strains compared with those of colistin monotherapy group.

**Conclusions:** LPC used as an adjuvant of colistin treatment may be helpful to reduce the severity and the resolution of the infection by MDR *A. baumannii*.

## INTRODUCTION

*Acinetobacter baumannii* is a gram-negative bacillus with high clinical relevance owing to the increase in the number of nosocomial infections caused by this pathogen, as well as its ability to develop resistance to most antimicrobial agents used by physicians (1). Treatment of *A. baumannii* infections, especially those caused by MDR strains is a major concern. In many areas of the world that have a high prevalence of MDR *A. baumannii*, few options of treatment are present and last resort treatments such as colistin are no longer effective in an increasing number of cases, leading to a 28-day mortality of 43% in hospitalized patients with bacteremia, ventilator-associated or hospital acquired pneumonia, or urosepsis (2). The number of antibiotics approved by the FDA cannot keep pace with the resistance mechanisms acquired by *A. baumannii*. Therefore, the development of new strategic antimicrobial therapeutic approaches, like the use of non-antibiotics in combination with one of the scarce but clinically relevant antibiotics, has become an urgent need.

A therapeutic alternative for infections by MDR *A. baumannii* is immune system modulation to improve the infection clearance. We previously successfully demonstrated the efficacy of lysophosphatidylcholine (LPC), a phospholipid involved in the recruitment and stimulation of immune cells (3–6) as a preemptive treatment in murine peritoneal sepsis and pneumonia experimental models by susceptible and MDR *A. baumannii* strains (7). Of note, LPC preemptive treatment in combination with colistin, tigecycline, or imipenem treatment has improved the *in vivo* antibacterial activity of these antimicrobials in murine experimental peritoneal sepsis and pneumonia by drug-susceptible and MDR *A. baumannii* (8). In the same line, LPC preemptive treatment in combination with ceftazidime has potentiated the *in vivo* antibacterial activity of ceftazidime in these severe infections models by MDR *Pseudomonas aeruginosa* (9). Recently, Yadav *et al*. have reported *in vitro* that LPC potentiate the effect of nonbactericidal concentration of polymexin B against the growth of *Pseudomonas aeruginosa* and *Klebsiella pneumoniae* (10).

Currently, there are no data regarding the efficacy of the direct treatment with LPC in combination with colistin against MDR *A. baumannii*. Therefore, the aim of this study was to evaluate the efficacy of the direct treatment with LPC in combination with colistin in murine experimental models of peritoneal sepsis and pneumonia by drug-susceptible and MDR clinical isolates of *A. baumannii*.

## MATERIALS AND METHODS

### Bacterial strains

Drug-susceptibe *A. baumannii* (Ab9) and MDR *A. baumannii* (Ab186) (resistant to imipenem, tigecycline, ciprofloxacin and ceftazidime) clinical strains were used in this study (8). Both strains were susceptible to colistin with MIC of 0.5 mg/L. The MIC of LPC against both strains was >8.000 mg/L (8). Ab9 was recovered from a wound surgical exudates, and Ab186 was recovered from blood cultures and belong to ST297 and ST2 (international clone II), respectively (8, 11).

### Antimicrobial agents and reagents

Clinical formulation of colistin methanesulfonate (Promixin®, Spain) was used. The anesthetic was 2:1 Ketamine hydrochloride® (Pfizer, Spain):Diazepam ® (Roche, Spain).

### Animals

Immunocompetent C57BL7/6 female mice weighing 18 to 20 g (Production and Experimentation Animal Center, University of Seville, Seville, Spain) were used. Animals were housed in regulation boxes and given free access to food and water. This study was carried out in strict accordance with the recommendations in the *Guide for the Care and Use of Laboratory Animals* (12). The protocol was approved by the Committee on the Ethics of Animal Experiments of the University Hospital of Virgen del Rocío of Seville, Spain (approval 1556-N-16).

### Experimental murine model of peritoneal sepsis

A previously characterized murine model of peritoneal sepsis caused by *A. baumannii* was used (8). Briefly, animals were inoculated intraperitoneally (ip.) with 0.5 mL of the 100% minimal lethal dose (MLD_100_) of the Ab9 (5.9 log_10_ CFU/mL) or Ab186 (5 log_10_ CFU/mL), mixed 1:1 with 10% porcine mucin (Sigma, Spain). LPC (Sigma, Spain) and colistin treatments were administered 4 h after bacterial inoculation. Groups of mice were randomly ascribed to the following groups: (i) controls (without treatment), (ii) LPC administered once i.p at 25 mg/kg 4 h after bacterial inoculation, (iii) colistin administered i.p at 20 mg/kg/8 h for 72 h (8) and (iv) the combination of colistin at 20 mg/kg/8 h with one dose of LPC at 25 mg/kg/d, and (v) the combination of colistin at 20 mg/kg/8 h with two doses of LPC at 25 mg/kg/d (first and second at 4 and 28 h, respectively, after bacterial infection).

Mortality was recorded over 72 h. After the death or the euthanization of the mice by sodium thiopental (Zambon S.p.A., Italy) at the end of the experiment period, aseptic thoracotomies were performed, and blood samples were obtained by cardiac puncture. Spleen and lungs were aseptically removed and homogenized (Stomacher 80®; Tekman Co.) in 2 mL of sterile 0.9% NaCl solution. Tenfold dilution of the homogenized spleen and lungs, and blood obtained by cardiac puncture, were plated onto sheep agar for the quantitative cultures (to determine the log_10_ CFU/g of spleen and lungs and log_10_ CFU/mL of blood).

### Experimental murine model of pneumonia

A previously described experimental murine pneumonia model was used to evaluate the efficacy of LPC as monotherapy and in combination with colistin against Ab9 and Ab186 strains (8). Briefly, the mice were anesthetized by 2:1 Ketamine hydrochloride:Diazepam, suspended vertically, and the trachea of each was then cannulated with a blunt-tipped metal needle. The feel of the needle tip against the tracheal cartilage confirmed the intratracheal location. A microliter syringe (Hamilton Co., Reno, NV) was used for the inoculation of 50 μL of bacterial suspension (10 and 9 log_10_ CFU/mL for Ab9 and Ab186 strains, respectively) which had been grown for 24 h in LB broth at 37°C and mixed at a 1:1 ratio with 0.9% NaCl solution containing 10% (wt/vol) porcine mucin. The mice remained in a vertical position for 3 min and then in a 30° position until they awakened. Treatment groups were similar to those for the experimental model of peritoneal sepsis. After death or sacrifice of the mice at the end of the experimental period, aseptic thoracotomies were performed, and blood was obtained by cardiac puncture and lungs were aseptically removed and homogenized. Quantitative data was obtained as described above to determine the log_10_ CFU/g of lungs and log_10_ CFU/mL of blood and mice mortality was recorded over 72 h.

### Statistical analysis

Group data are presented as means ± standard errors of the means (SEM). Differences in the bacterial spleen, lung and blood concentrations (mean ± SEM log CFU per gram of tissue or per ml of blood) were assessed by analysis of variance (ANOVA) and the post hoc Dunnnett test Differences in mortality (%) and blood sterility (%) between groups were compared by χ^2^ test. *P* values of <0.05 were considered significant. The SPSS (version 23.0; SPSS Inc.) statistical package was used.

## RESULTS

### Efficacy of LPC in combination with colistin in murine experimental model of peritoneal sepsis

The efficacies of colistin and LPC in monotherapies and in combination against Ab9 and Ab186, expressed as survival and bacterial concentrations in spleen, lungs and blood, are shown in the tables 1 and 2.

**Table 1.** Therapeutic effect of one or two doses of LPC in combination with colistin in murine model of peritoneal sepsis with *A. baumannii* Ab9.

| Treatment | *n* | Spleen | Lung | Blood | Mortality |
| --- | --- | --- | --- | --- | --- |
|  |  | (log<sub>10</sub> CFU/g) | (log<sub>10</sub> CFU/g) | (log<sub>10</sub> CFU/ml) | (%) |
| CTL | 10 | 9.55 ± 0.09 | 9.85 ± 0.72 | 8.59 ± 0.04 | 100 |
| LPC | 8 | 9.82 ± 0.08 | 9.69 ± 0.91 | 9.20 ± 0.04<sup>a</sup> | 100 |
| CST | 8 | 4.48 ± 0.30<sup>a,b</sup> | 4.17 ± 0.29<sup>a</sup> | 3.26 ± 0.40<sup>a,b</sup> | 25<sup>a</sup> |
| LPC1 + CST | 8 | 3.98 ± 0.66<sup>a,b</sup> | 3.83 ± 0.65<sup>a</sup> | 2.92 ± 0.58<sup>a,b</sup> | 0<sup>a</sup> |
| LPC2 + CST | 8 | 3.42 ± 0.50<sup>a,b</sup> | 3.13 ± 0.46<sup>a</sup> | 1.85 ± 0.38<sup>a,b</sup> | 0<sup>a</sup> |

CTL, control (no treatment); LPC, lysophosphatidylcholine; CST, colistin; LPC1, one dose of lysophosphatidylcholine; LPC2 two doses of lysophosphatidylcholine.
<sup>a</sup> *P*<0.05 compared to the controls
<sup>b</sup> *P*<0.05 compared to LPC group

**Table 2.** Therapeutic effect of one or two doses of LPC in combination with colistin in murine model of peritoneal sepsis with *A. baumannii* Ab186.

|  |  | **Spleen** | **Lung** | **Blood** | **Mortality** |
| --- | --- | --- | --- | --- | --- |
| **Treatment** | **n** | **(log<sub>10</sub> CFU/g)** | **(log<sub>10</sub> CFU/g)** | **(log<sub>10</sub> CFU/ml)** | **(%)** |
| **CTL** | 13 | 9.79 ± 0.06 | 9.63 ± 0.13 | 8.89 ± 0.03 | 100 |
| **LPC** | 8 | 10.48 ± 0.03 | 10.03 ± 0.03 | 9.29 ± 0.03 | 100 |
| **CST** | 8 | 2.86 ± 1.54<sup>a,b</sup> | 2.90 ± 1.57<sup>a,b</sup> | 2.19 ± 1.63<sup>a,b</sup> | 75 |
| **LPC1 + CST** | 12 | 1.58 ± 0.48<sup>a,b</sup> | 1.43 ± 0.54<sup>a,b</sup> | 0.22 ± 0.21<sup>a,b</sup> | 0<sup>a,b</sup> |
| **LPC2 + CST** | 12 | 0.22 ± 0.29<sup>a,b</sup> | 0.75 ± 0.32<sup>a,b</sup> | 0.08 ± 0.12<sup>a,b</sup> | 0<sup>a,b</sup> |

CTL, control (no treatment); LPC, lysophosphatidylcholine; CST, colistin; LPC1, one dose of lysophosphatidylcholine; LPC2 two doses of lysophosphatidylcholine.
<sup>a</sup> *P*<0.05 compared to the controls
<sup>b</sup> *P*<0.05 compared to LPC group

#### (i) Survival

Tables 1 and 2 show that colistin alone and in combination with one and two doses of LPC increased mice survival compared with that of the control group for Ab9 and Ab186 (*P*<0.05). In contrast, LPC in monotherapy did not reduce mice mortality.

#### (ii) Bacterial clearance from spleen, lungs and blood

Tables 1 and 2 show that monotherapy with colistin cleared Ab9 and Ab186 from the spleen, lungs and blood by 5.07 and 5.68 log_10_ CFU/g, and 5.33 log_10_ CFU/mL (*P*<0.05; Ab9), respectively, and 6.93 and 6.73 CFU/g, 6.7 log_10_ CFU/mL (*P*<0.05; Ab186), respectively, compared with the levels of the control group. One dose of LPC in combination with colistin decreased spleen, lungs and blood concentrations of Ab9 and Ab186 by 5.57 and 6.02 log_10_ CFU/g, and 5.67 log_10_ CFU/mL (*P*<0.05; Ab9) respectively, and 8.21 and 8.2 log_10_ CFU/g, and 8.67 log_10_ CFU/mL (*P*<0.05; Ab186), respectively, compared with the levels for the control group. In addition, the increase of the dose of LPC has slightly increased the bacterial clearance. Two doses of LPC in combination with colistin reduced the bacterial burden in spleen, lungs and blood by 6.13 and 6.72 log_10_ CFU/g, and 6.74 log_10_ CFU/mL (*P*<0.05; Ab9), respectively, and 9.57 and 8.88 log_10_ CFU/g, and 8.81 CFU/mL (*P*<0.05; Ab186), respectively, compared with the levels for the control group. Of note, one dose of LPC in combination with colistin decreased spleen, lungs and blood concentrations of Ab9 and Ab186 by 5.84 and 5.86 log_10_ CFU/g, and 6.28 log_10_ CFU/mL, respectively (*P*<0.05; Ab9), and 8.9 and 8.6 log_10_ CFU/g, and 9.07 log_10_ CFU/mL (*P*<0.05; Ab186), respectively, compared with the levels for the LPC monotherapy group. Finally, two doses of LPC in combination with colistin decreased spleen, lungs and blood concentrations for Ab9 and Ab186 by 6.4 and 6.56 log_10_ CFU/g, and 6.74 log_10_ CFU/mL (*P*<0.05; Ab9), respectively, and 10.26 and 9.28 log_10_CFU/mL, and 9.21 log CFU/mL (*P*<0.05; Ab186), respectively, when compared with the levels for the LPC monotherapy.

### Efficacy of LPC in combination with colistin in murine experimental model of pneumonia

The efficacies of colistin and LPC in monotherapies and in combination against Ab9 and Ab186, expressed as survival and bacterial concentrations in spleen, lungs and blood, are shown in the tables 3 and 4.

**Table 3.** Therapeutic effect of one or two doses of LPC in combination with colistin in murine pneumonia model with *A. baumannii* Ab9.

| Treatment | *n* | Lung | Blood | Mortality |
| --- | --- | --- | --- | --- |
|  |  | (log<sub>10</sub> CFU/g) | (log<sub>10</sub> CFU/ml) | (%) |
| CTL | 8 | 9.64 ± 0.55 | 7.95 ± 0.83 | 87.5 |
| LPC | 8 | 9.13 ± 0.28 | 7.27 ± 0.04 | 100 |
| CST | 8 | 3.11 ± 1.18<sup>a,b</sup> | 2.14 ± 0.57<sup>a,b</sup> | 12.5<sup>a,b</sup> |
| LPC1 + CST | 8 | 2.88 ± 1.12<sup>a,b</sup> | 1.87 ± 0.6<sup>a,b</sup> | 12.5<sup>a,b</sup> |
| LPC2 + CST | 8 | 1.90 ± 1.13<sup>a,b</sup> | 1.31 ± 0.80<sup>a,b</sup> | 12.5<sup>a,b</sup> |

CTL, control (no treatment); LPC, lysophosphatidylcholine; CST, colistin; LPC1, one dose of lysophosphatidylcholine; LPC2 two doses of lysophosphatidylcholine.
<sup>a</sup> *P*<0.05 compared to the controls
<sup>b</sup> *P*<0.05 compared to LPC group

**Table 4.** Therapeutic effect of one or two doses of LPC in combination with colistin in a murine pneumonia model with *A. baumannii* Ab186.

| Treatment | *n* | Lung | Blood | Mortality |
| --- | --- | --- | --- | --- |
|  |  | (log<sub>10</sub> CFU/g) | (log<sub>10</sub> CFU/ml) | (%) |
| CTL | 8 | 9.21 ± 0.45 | 7.76 ± 0.39 | 100 |
| LPC | 8 | 9.01 ± 0.07 | 7.33 ± 0.03 | 100 |
| CST | 8 | 1.66 ± 0.49<sup>a,b</sup> | 0.97 ± 0.30<sup>a,b</sup> | 0<sup>a,b</sup> |
| LPC1 + CST | 8 | 1.11 ± 0.54<sup>a,b</sup> | 0.59 ± 0.29<sup>a,b</sup> | 0<sup>a,b</sup> |
| LPC2 + CST | 8 | 0.65 ± 0.43<sup>a,b</sup> | 0.43 ± 0.28<sup>a,b</sup> | 0<sup>a,b</sup> |

CTL, control (no treatment); LPC, lysophosphatidylcholine; CST, colistin; LPC1, one dose of lysophosphatidylcholine; LPC2 two doses of lysophosphatidylcholine.
<sup>a</sup> *P*<0.05 compared to the controls
<sup>b</sup> *P*<0.05 compared to LPC group.

#### (i) Survival

Tables 3 and 4 show that colistin alone and in combination with one and two doses of LPC increased mice survival compared with that of the control group for Ab9 and Ab186 (*P*<0.05). In contrast, LPC in monotherapy did not reduce mice mortality.

#### (ii) Bacterial clearance of lungs and blood

Tables 3 and 4 show that monotherapy with colistin cleared Ab9 and Ab186 from the lungs and blood by 6.53 and 5.81 log_10_ CFU/g and mL (*P*<0.05; Ab9), respectively, and 7.75 and 6.79 log_10_ CFU/g and mL (*P*<0.05; Ab186), respectively, compared with the levels of the control group. One dose of LPC in combination with colistin decreased lungs and blood concentrations of Ab9 and Ab186 by 6.76 and 6.08 log_10_ CFU/g and mL (*P*<0.05; Ab9) respectively, and 8.1 and 7.17 log_10_ CFU/g and mL (*P*<0.05; Ab186), respectively, compared with the levels for the control group. In addition, the increase of the dose of LPC has slightly increased the bacterial clearance. Two doses of LPC in combination with colistin reduced the bacterial burden in lungs and blood by 7.74 and 6.64 log_10_ CFU/g and mL (*P*<0.05; Ab9), respectively, and 8.56 and 7.33 CFU/g and mL (*P*<0.05; Ab186), respectively, compared with the levels for the control group.

Finally, one and two doses of LPC in combination with colistin decreased the lungs concentrations of Ab9 by 6.25 and 7.25 log_10_ CFU/g (*P*<0.05) and Ab186 by 7.9 and 8.36 log_10_ CFU/g (*P*<0.05), compared with the levels for the LPC monotherapy. Similar results were observed in blood with a reduction of 5.4 and 5.95 log_10_ CFU/mL (*P*<0.05; Ab9), and 6.74 and 6.9 log_10_ CFU/mL (*P*<0.05; Ab186) compared with the levels for the LPC monotherapy.

## DISCUSSION

Previous studies from our group demonstrated that preemptive LPC monotherapy and in combination with antibiotics such as colistin reduced bacterial tissues loads and bacteremia and increased mice survival in murine experimental models of severe infections by *A. baumannii* (7, 8). Even though LPC as preemptive monotherapy and in combination with colistin presented remarkable results, we hypothesized that it may be given as direct treatment in combination with colistin.

Currently, colistin is among the last treatments available worldwide, being a last resort against MDR *A. baumannii* strains. Nevertheless, its therapeutic efficacy using optimal doses is limited, being effective just in the 60% of patients infected with a MDR strain susceptible to colistin (13, 14). For that reason, two different clinical isolates have been chosen, one drug-susceptible and one MDR, both susceptible to colistin. In the present study, monotherapy with colistin against drug-susceptible and MDR *A. baumannni* strains significantly reduced bacterial concentrations in spleen, lungs and blood and increased mice survival comparing with the control group. However, it is important to highlight that colistin monotherapy presented a mortality rate of 75% in the case of the MDR strain in the peritoneal sepsis model. This result revealed a failure in the treatment with colistin, and the mice survival values are similar and even higher to the rates obtained in the clinical practice when dealing with a colistin-susceptible strain with highly resistant pattern. Accordingly with our hypothesis, treatment with one or two doses of LPC in combination with colistin in a peritoneal sepsis model increased (without statistical difference) mice survival and reduced bacterial loads in tissues and blood, comparing with colistin monotherapy. No differences were found between a single dose and multiple doses of LPC. It is noteworthy to mention that higher efficacy of the combination LPC plus colistin was observed against the MDR strain Ab186, where survival rates were markedly increased. In the case of the pneumonia model, no differences were found in survival rates comparing with colistin monotherapy but a decrease in lungs and blood bacterial concentrations were observed.

Differences in bacterial concentrations were not due to different pharmacokinetic parameters between strains, since the MIC value of colistin for both strains is 0.5 mg/l. Different response to the colistin treatment may be explained by immune responses caused by both strains. Indeed, Ab9 induced more TNF-alpha release than that of the Ab186 (8). Other studies reported by our group showed that a drug-susceptible *A. baumannii* strain induced more TNF-α and interleukin 6 releases than MDR and pan-drug resistant *A. baumannii* clinical isolates (15, 16). In line with this hypothesis, increased lethality and severity of the infection by *A. baumannii* was observed when neutrophils are depleted, together with a delayed production of cytokines involved in neutrophil function such as TNF-α, interleukin 1, keratinocyte chemoattractant protein (KC/CXCL1) and macrophage inflammatory protein (MIP-1) (17). Neutrophils are essential players during *A. baumannii* infection and present an important role against sepsis and pneumonia infection (18, 19). It was reported that LPC blocks neutrophil deactivation during murine cecal ligation and puncture model as well as increased the bactericidal activity of these immune cells (20). Thus, the additive action of LPC to the antibiotic treatment may be due to enhanced activity of neutrophils.

Interestingly, direct treatment with LPC in combination with colistin presents similar efficacy than preemptive treatment with LPC in combination with colistin against MDR Ab186 strain. A reduction of the bacterial burden in spleen and lungs around 2 log_10_ CFU/g in a murine model of peritoneal sepsis and pneumonia models comparing with the LPC in combination with colistin preemptive treatment was observed (8). This comparison increases the interest towards LPC as a future adjuvant therapy with colistin, which may reduce the apparition of resistance to antibiotics (21).

In summary, the present study suggests that direct treatment with LPC in combination with colistin improves the *in vivo* antibacterial activity in murine experimental models of peritoneal sepsis and pneumonia by MDR *A. baumannii*.

## ACKNOWLEDGEMENTS

This study was supported by the Instituto de Salud Carlos III, Proyectos de Investigación en Salud (grants PI13/01744, PI16/01306 and PI16/01378) and by Plan Nacional de I+D+i 2013-2016 and Instituto de Salud Carlos III, Subdirección General de Redes y Centros de Investigación Cooperativa, Ministerio de Ciencia, Inno-vación y Universidades, Spanish Network for Research in Infectious Diseases (REIPI RD16/0016/0009) –co-financed by European Development Regional Fund “A way to achieve Europe”, Operative program Intelligent Growth 2014-2020. Younes Smani is supported by the Subprograma Miguel Servet Tipo I from the Ministerio de Economía y Competitividad of Spain (CP15/01358).

